# A Thermostable Tetanus/diphtheria (Td) Vaccine in the Stablevax^™^ Pre-Filled Delivery System

**DOI:** 10.1101/2023.02.16.526924

**Authors:** Juana de la Torre Arrieta, Daniela Briceño, Ivan Garcia de Castro, Bruce Roser

**Affiliations:** Stablepharma Ltd 4 Queen Square Bath BA1 2HA Somerset U.K.

**Author notes:** This research did not receive any specific grant from funding agencies in the public, commercial, or not-for-profit sectors.

**Keywords:** accelerated-ageing, fridge-free, pre-filled syringe, reticulated-sponge, stable-vaccine, thermostable, trehalose

## Abstract

A pre-filled syringe for the long-term, room-temperature storage and injection of vaccines is described. Stabilisation was achieved by drying from a trehalose-containing buffer which formed an inert soluble glass distributed in the internal interconnected voids in a compliant reticulated medical-grade sponge. The sponge is stored inside the barrel of the syringe and the vaccines are re-solubilised by the aspiration of water. The syringe contains the sponge throughout the filling and drying processes in manufacture, and in transport, stockpiling and finally injection. The active vaccine is delivered to the patient in the normal injection process by depressing the plunger, which compresses the sponge to completely expel the dose. There was full recovery of vaccine potency, after 7-10 months @ 45°C, as shown by complete protection against active toxins in immunised Guinea pigs.

## INTRODUCTION

Vaccines require continuous refrigeration, from factory to patient in the “cold chain”, which greatly complicates both their worldwide distribution and stockpiling. This is especially acute with modern nucleic acid-based vaccines which, in the SARS Cov-2 pandemic, have proved to be the most efficacious but are also the most fragile. These require freezing for stability, and in the case of the Pfizer/BioNTech mRNA vaccines for example, were recommended ultra-low temperatures of –60 to –90°C [1]

However, even the least demanding refrigerated storage between 2-8°C, which is required for most other vaccines, is not universally available, or entirely reliable [2]. Breakdowns in correct temperature storage result in about 50% of all vaccines being damaged by heating [3] or, by freezing [4].The incidence of accidental freezing of sensitive vaccines, reviewed by Matthias *et al* [5] noted that in a total of 2,236 stored adjuvanted vaccines reported in 41 publications, over 40% had been accidentally frozen.

What is urgently needed in this globally connected World, in order to pre-empt the transmission of new pathogens or new waves of mutants, is that vaccination can be quickly extended in a pandemic to everyone Worldwide. This will require a technology to produce effective safe vaccines that can be manufactured and distributed quickly [6–8] and are resistant to adverse storage or transportation conditions. An ideal solution to this problem would be a completely thermostable formulation of vaccines pre-packed as single doses in the injection syringe itself [9]. This would also reduce costs, since other equipment required by current vaccines such as vials, sterile empty syringes and the low temperature storage and monitoring equipment required for the cold chain, would all be unnecessary.

A wide range of pharmaceuticals can be thermally stabilized by drying in soluble glasses, particularly sugar glasses [10–12], and most effectively trehalose [13–15]. These dry, stabilized actives are unaffected by both high and freezing temperatures. A most important mechanism underlying the remarkable stabilization of molecules by sugars is the ability of drying solutions to undergo glass-transformation rather than crystallisation. Trehalose, unlike most other sugars is chemically very inert and so cannot react with and damage the product. It also readily forms stable glasses [16].

Pharmaceuticals currently stabilised by drying are usually freeze-dried. A correctly processed “elegant” freeze-dried plug is highly valued [17], largely because of its uniform porosity, which facilitates reconstitution of the dried product by avid absorption of water and rapid dissolution. StablevaX technology uses a uniformly porous sponge that mimics this property. It also facilitates manufacturing by absorbing the liquid vaccine containing stabilisers in a widely dispersed distribution with a very large surface area for water evaporation and rapid drying at higher temperatures. For final administration to the patient, the aspiration of water into the syringe starts the dissolution of the soluble sugar glass containing the vaccine. After the product is dissolved, and during injection, the plunger fully compresses the sponge, which actively expels the contained liquid, and the complete dose of the pharmaceutical is delivered.

We have previously described and patented a method, using trehalose-based stabilisation, for the implementation of this new technology [9]. We now report the results of a project (codename STVX-01) to develop the first “fridge Free” commercial product by applying a further development of the technology to a widely used, thermally unstable, adjuvanted, divalent vaccine. Since administering vaccinations require the use of a new sterile syringe for each patient, the StablevaX process is designed to use the syringe itself as the sole vaccine container throughout the manufacturing steps of filling with vaccine, drying and packaging, and then its transportation, stockpiling, and storage, and finally, its injection.

Considering the issues of freedom to operate, commercial availability, market need, and a formulation that would be a stringent test of the StablevaX process, the first candidate chosen for the development of STVX-01 was a commercial Tetanus diphtheria (Td) vaccine, (Tetadif, BulBio BB-NCIPD Ltd) [18]. While diphtheria and tetanus toxoids are among the most stable toxoids in commonly used vaccines, they are only stable for years if continuously refrigerated at 2-8°C. At 37°C they are stable for 2-6 months, and at 45°C, for 2-4 weeks, and are destroyed within 3-5 hours at >60°C [19].

Adjuvants are essential to ensure optimal immune responses to many vaccines [20], but aluminium hydroxide-adjuvanted vaccines are very susceptible to drying and freezing damage since the highly hydrated structure of the adjuvant gel is irreversibly damaged and forms insoluble aggregates [21]. This Td divalent vaccine, adjuvanted with aluminium hydroxide, was therefore chosen to provide a demonstration that with StablevaX, both the soluble antigens and the particulate adjuvant component of the vaccine could be successfully dried and quantitatively recovered in a fully functional state even after prolonged high-temperature storage.

## MATERIALS AND METHODS

### Preparation of sponges and syringes

15 mm thick sheets were sliced from commercial reticulated polyurethane foam Buns (#SPU6. UFP Technologies, Newburyport, Mass. USA) with pore size of 80 pores per inch. Cylindrical sponges of 8 mm diameter (volume 754 mm^3^) were cut with die punches (AQF Medical, Co. Meath, Ireland). These sponges were washed with water at 80°C containing 0.1% polysorbate 80, then rinsed twice in deionised water and dried at 40°C overnight in a convection oven (Digitronic, J.P. Selecta^®^, Barcelona, Spain). To minimise any residual particles, a second washing step which included sonication, was used (Ultrasons, J.P. Selecta^®^, Barcelona, Spain) and the sponges were again dried at 40°C overnight. Clean sponges were then autoclaved at 120°C for 15 minutes (Presoclave II, J.P. Selecta^®^, Barcelona, Spain) and loaded into sterile 3ml syringes (Plastipak TM 3ml Syringe Luer-Lok™ Tip, Becton Dickinson, USA).

### Vaccine Compounding

For pilot scale manufacturing of batches of 35 STVX-01 syringes, 50 doses of vaccine were prepared to allow for wastage. The compounded vaccine consisted of bulk vaccine (Tetadif, BulBio, Sofia, Bulgaria) to which was added a filter-sterilized 60% w/v solution of crystalline trehalose dihydrate (Pfanstiehl GmbH, Zug, Switzerland) and Polysorbate 80 to provide a final concentration of 0.243 M trehalose and 0.1% v/v Polysorbate 80.

### Sponge loading

Each sponge was filled with 0.65 ml of the compounded vaccine via the nozzle of the syringe by using a repeater pipette (Multipette^®^ E3/E3x, Eppendorf^®^, Hamburg, Germany) with the disposable plastic tip cut to make a press-fit seal with the hub of a 23-gauge hypodermic needle. Once the compounded vaccine was delivered into the barrel of the syringe and absorbed by the sponge, any air bubbles trapped in the sponge were removed and any stray vaccine droplets from the walls of the syringe were delivered to the sponge, by limited movements of the plunger and gentle tapping of the syringes.

### Drying Rig

“In-syringe” drying was achieved in a rig fabricated in Stainless Steel. This rig located 35 syringes in an array of 7 rows of 5, at 17 mm centres. Drying air was delivered from a heated manifold by narrow stainless tubes in alignment with the syringe nozzles and raised and lowered on polished SS guide rods. Temperature was monitored of the incoming air as well as in the core of the sponge with a KPS-TM340 2-channel digital thermometer and 2 thermocouples. To establish the best drying conditions, the evaporation of water was followed by monitoring the weight loss of single StablevaX units using different air flow rates, from 2 Litres per Minute (L/m) to 10 L/m, and an air temperature of 42°C. Weights were recorded at intervals on an electronic balance (Mettler Toledo, XS105, Barcelona, Spain). The final drying conditions used for the functional evaluations of the product were overnight rig drying with 42°C air temperature followed by secondary drying at 40°C for 4hr in a vacuum oven (Binder, Tuttlingen, Germany) with vacuum of 0.1 mbar.

This drying protocol including secondary drying was used to ensure complete dryness for subsequent stability testing. For commercial production it is highly likely that higher drying temperatures will result in a much faster single step process.

### Residual Water content

Moisture was measured by Karl Fisher titration using a KF1000 Titrator (Hach Lange, GMBH, Dusseldorf, Germany).

### Residual Particles

A microscopic count, (USP788 Particulate matter in injections - Method B) was used to evaluate the presence of residual particles. Because of the high concentration of adjuvant particles in the vaccine itself, any possible contamination of the injection with foreign particles from the sponge was checked using mock formulations dried in sponges containing the buffers and excipients minus the adjuvanted vaccine. Ten syringes were each rehydrated with 2.5 ml of particle-free water. The eluents were pooled and collected by filtration on to a 0.2 μm cellulose nitrate filter, (Sartorius Stedim biotech Gmbh, Goettingen, Germany), dried and examined in a binocular microscope using a measuring reticle (Nexius Zoom, Euromex, Arnhem, Holland). Freedom from accidental release of particles during use was also verified following deliberate compression of dry sponges by depression of the plunger in syringes before rehydration such as could happen during transportation or warehousing or by misuse.

### Cytotoxicity

Clean foam cylinders placed in BD 3ml syringes, as described in “**Preparation of sponges and syringes**” (above), were also sterilised by electron beam radiation at 50kGy (Ionisos Iberica S.L. Cuenca, Spain) and were checked for absence of cytotoxicity (Biolab S.L. Madrid, Spain), by exposing tissue culture media extracts of the sponges to *in vitro* cell cultures. Cytotoxicity was evaluated according to the ISO Guidelines, UNE-EN ISO. 10993-5:2009. Biological evaluation of medical devices. Part 5: Test for in vitro cytotoxicity, and UNE-EN ISO 10993-12:2013. Biological evaluation of medical devices. Part 12: Sample preparation and reference materials. The extracts obtained were sterilised by 0.22 μm filtration and diluted to obtain concentrations of 100, 75, 50 and 25 % extract. Cell line CCL 81 “Vero” cells were incubated with the extracts for 24 h and cell viability was determined by MTT (3-(4,5-Dimethylthiazol-2-yl)-2,5-diphenyltetrazolium bromide) assay.

### Reconstitution of Vaccine

To allow for the retained void volume of ~ 100 μl, 0.60 ml of water for injection was aspirated into the syringe. To ensure full rehydration and homogeneity in the resuspension, it was necessary to flick the filled syringe vigorously for 60 seconds (approx. 60 times). This ensured that any trapped air bubbles were released, and the rehydrating liquid had completely saturated the sponge.

### Adjuvant Release

Two methods were used to measure recovery of Al(OH)_3_ adjuvant from the sponge. **Method A** measured particulate Al(OH)_3_ by Absorbance at 960 nm by a modification of the method of Lai et al. [22] in Quartz SUPRASIL® 10mm pathlength cuvettes in a UV-1800 UV spectrophotometer, (Shimadzu Corporation, Kyoto Japan). The relationship between the adjuvant concentration and absorbance was shown to be linear using eight concentrations of Alhydrogel® (Croda Iberica SA Barcelona, Spain) from 0.02 to 2.5 mg/ml. The measurements were checked by comparison with the documented values in 4 batches of commercial Td-vaccine, which showed close correspondence.

**Method B** was the chemical estimate of aluminium using European Pharmacopoeia monograph Ph.Eur 2.5.13. “Aluminium in Adsorbed Vaccines” and was always employed for final formulation candidates and pilot production batches.

### Adjuvant gel structure integrity

1 x G Sedimentation columns were used to verify that the native gelatinous structure of the aluminium hydroxide is retained after freezing or drying. 10 test samples were resuspended to the original vaccine volume and pooled. 4 ml of the pooled samples were aspirated into 5ml plastic pipettes, sealed at the bottom, and left standing vertically in a clamp. After 24h the height of the column was measured and compared with the sedimentation column generated by 4ml of the same batch of Tetadif liquid vaccine as a positive control.

### ELISA assays

Recovery of Tetanus (T) and Diphtheria (d) toxoids were measured after desorption from the Al(OH)_3_ adjuvant, by Enzyme Linked Immunosorbent Assay (ELISA) against biological standards from National Institute for Biological Standards and Control (NIBSC, South Mimms, Hertfordshire, UK) [23,24] After noting problems in detecting the Tetanus antigen content when it was desorbed using the citrate method, and following the recommendation from the Division of Bacteriology at NIBSC, tetanus antigen samples were prepared by desorption with buffered Ethylenediaminetetraacetic acid (EDTA) (Sigma-Aldrich Chemie GmbH Taufkirchen Germany). Diphtheria antigen samples were prepared by desorption with sodium citrate (Sigma-Aldrich Chemie GmbH Taufkirchen, Germany) [23]. Each antigen concentration was determined by UV absorbance at 450 nm in a plate reader (iMark™ Microplate Absorbance Reader, BioRad, Japan).

### Adjuvant: Electronic Particle-size Measurement

Samples were measured by dynamic light scattering (Zeta sizer Nano ZS, Malvern Panalytical Ltd, Malvern U.K.) using dual wavelengths: 466 nm and 633nm. The test samples were diluted two-fold with particle free distilled water immediately before recording the values. For each sample, an average of three separate readings was recorded. The data analysis was performed using Zetasizer Software Version 7.01 (Malvern Panalytical Ltd.).

### In Vivo potency assay (Lethal challenge)

Following Ph. Eur. 2.7.8 “Assay of Tetanus Vaccine Adsorbed” and using an internal BulBio minor modification, the potency of the protection against disease given by STVX-01 was determined by administering, to batches of 8 healthy guinea-pigs, a dose of 0.5 ml from a 4 step 2-fold dilution series of either the fresh Tetadif liquid vaccine or the stabilised STVX-01 vaccine that had been stored at 45°C for 7-10 months. After 28 days, all immunised animals were challenged subcutaneously with 1 ml of an active toxin solution containing 50 times the 50% paralytic dose (for Tetanus) or 100 times the 50% lethal dose (for Diphtheria). Challenged animals were observed daily for 5 days post challenge and any dead animals were removed and counted. All protection studies were calibrated with internal House-Standard vaccines which were periodically checked against NIBSC official standards. We report here potency data after 7–10-month storage at 45°C for Diphtheria and tetanus respectively. Potency of the test material relative to the potency of the standard preparation was calculated on the basis of the “number of survivors / number of challenged” ratios. The results were calculated by the CombiStats Programme (EDQM, Council of Europe).

## RESULTS

### Characterisation of components

#### Syringe: Requirements

- Three-component design to allow the elastomer plunger seal to “rest” half-way up the barrel.
- 2.5-3 ml volume to accommodate the sponge and rehydration procedure.
- Low void volume.
- Transparent to allow viewing of contents.
- Supplied non-sterile and disassembled allowing for insertion of the cleaned sponge prior to assembly and sterilization.
- Sterilizable by standard means (radiation, ethylene oxide or autoclave).

The widely used standard Beckton Dickinson (“BD”) 3 ml polypropylene syringes, and Terumo 2.5 ml syringes were used in this research.

#### Sponge: Requirements

The major industrial and medical use for porous sponges is the filtration of gaseous or liquid streams for removing particles. Since we needed a sponge with the opposite functionality, i.e. to completely release a particulate vaccine into a liquid stream, an extensive examination of available porous matrices was undertaken to find the best materials. The desirable criteria for the search were deemed to be:

- non-toxic.
- Not “adhesive” electrostatically or chemically for antigens or adjuvants.
- Shed minimal particles or leachables.
- Correct pore size with high and interconnected porosity.
- Easily cut to shape.
- Sterilizable by autoclave, radiation, or ethylene oxide.
- Compressible and resilient.
- Minimal swelling in water.
- Structurally stable without becoming brittle over time.
- Inexpensive.

Over 50 sponges made from foam materials from multiple manufacturers were tested for their capacity to absorb 0.65 ml of compounded vaccine and hold it all *in situ* by capillarity while being dried. Those passing this test were then used for recovery studies examining their ability to deliver the full dose of adjuvant with its gel structure unchanged, and the original quantity of intact adjuvant-bound antigen. Sponges selected for detailed evaluation were open cell sponges of three main chemical types, reticulated polyurethane, cellulose, or reticulated melamine formaldehyde. Final evaluation by examining particle release, swelling and shrinkage, and elutable contaminants, narrowed the choice to certain reticulated polyurethane sponges (which also had the advantage of having already been being clinically approved as dressings for open wounds)

#### Particle release

Microscope analysis showed there was an average of 220 particles between 10 and 150 μm and 0 particles (fibres) >500 μm, well within acceptable limits of 3,000 and 300 particles of those sizes respectively.

#### Cytotoxicity

Absence of cytotoxicity was established according to the ISO Guidelines by adding various concentrations of the extracts (100%, 50%, 33%, and 25%), to Vero cell cultures. Sponge cylinders were demonstrated to be non-cytotoxic as the test met the acceptance criteria of a cell viability better than 70% in all concentrations tested,

### Laboratory Scale Manufacturing Process

#### Sponge loading

Sterile syringes were disassembled, and one sterile empty sponge was inserted into each barrel. Syringes were then reassembled and positioned vertically to fill each one with 0.65 ml of compounded vaccine. This was delivered using a repeater pipette through the syringe nozzle via a 23-gauge hypodermic needle which was attached to its modified disposable plastic tip which had been cut to make a press-fit seal with the needle hub. After loading, any air bubbles trapped in the sponge could be removed and any stray vaccine droplets on the walls of the syringe absorbed into the sponge by gently moving the plunger and tapping the syringe. Over 30 batches of 35 x STVX-01 syringes were filled and dried in a custom rig in this way.

#### Air Flow

An enlarged view of the loaded sponge in the syringe barrel shows the flow of drying air (red arrows) around the sponge **F** which then exits **G** to atmosphere via the nozzle by flowing beside the air tube.

Preliminary studies using a single STVX-01 syringe equipped with thermistors to measure input dry air temperature and the temperature within the loaded sponge were used to provide evidence that this approach was valid. Drying of the sponge was monitored by weighing the syringe and contents at intervals through the drying process. To minimise possible damage to the drying vaccine, a low temperature, high flow rate, regime of airflow was chosen. The temperature of the drying air was fixed at 42°C and drying runs were conducted at vigorous flow rates from 2 to 10 L/m to define suitable conditions for production of the batches for functional studies.

10 Litres per minute was found to be too violent causing chaotic movement of the sponge and splashing on the barrel walls. Flow rates of 6, 4, and 2 L/m were smooth and easily controlled.

As expected, the rate of drying was proportional to flow rate and was found to closely follow an exponential function Figure 2.

**Figure 1.**
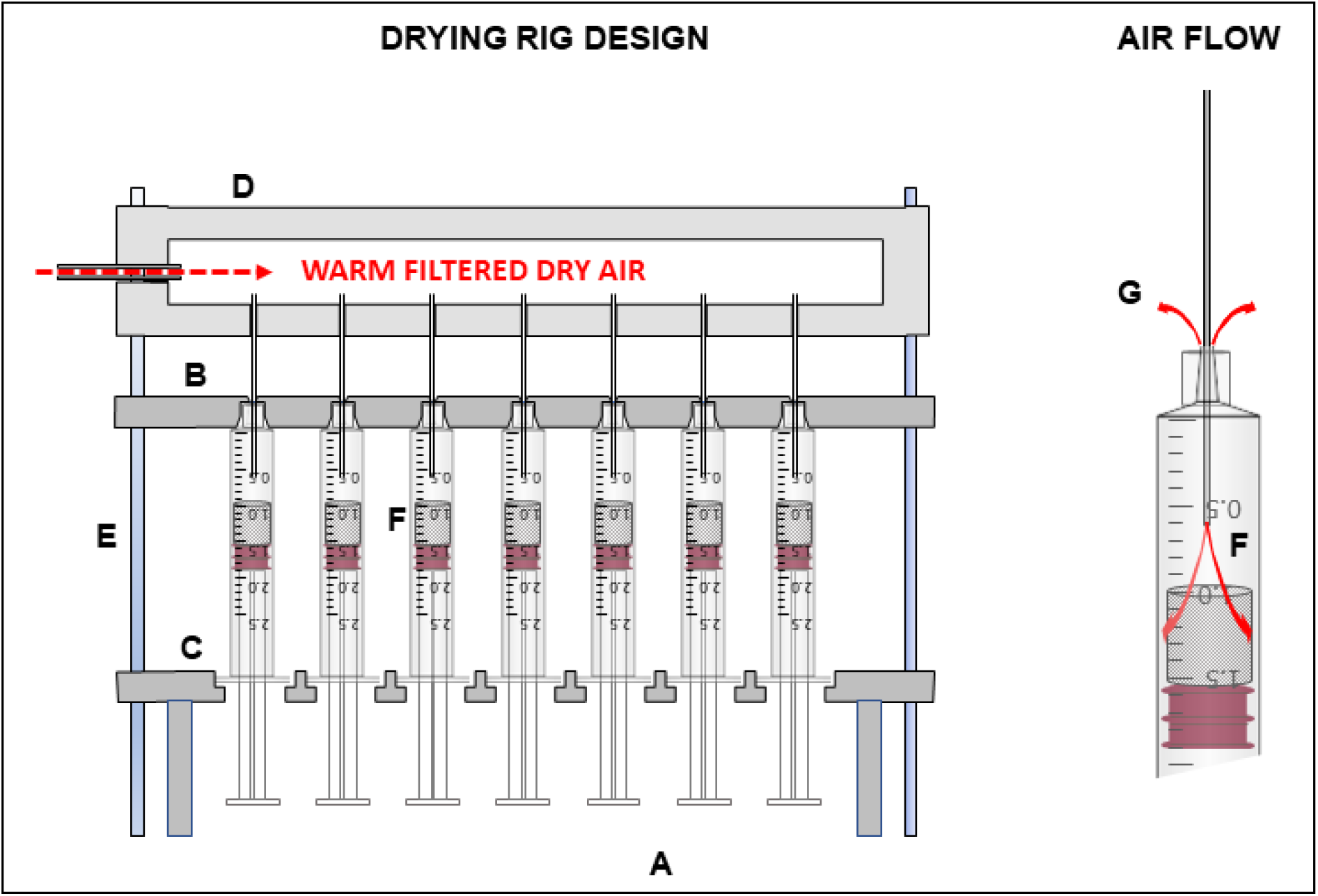
Drying rig design. The rig consisted of two parts. An assembly **A** to precisely locate 35 syringes containing the sponges loaded with compounded vaccine in a 7 x 5 array at 17 mm centres, and a drying air plenum **D**. The syringes are firmly held between two locator plates **B** & **C** with 35 holes machined to fit the syringe nozzle **B** and the barrel flange **C** respectively. The bottom locator plate **C** sits on 4 integral corner legs. Both locator plates **B**& **C** were drilled in the corners to accommodate 4, 10 x 0.5 cm polished Stainless rods **E** fixed to the corners of the air plenum **D** which precisely align the drying tubes of the plenum with the syringe nozzles and regulate the depth of their penetration into the syringe barrels as they are lowered into position.

**Figure 2.**
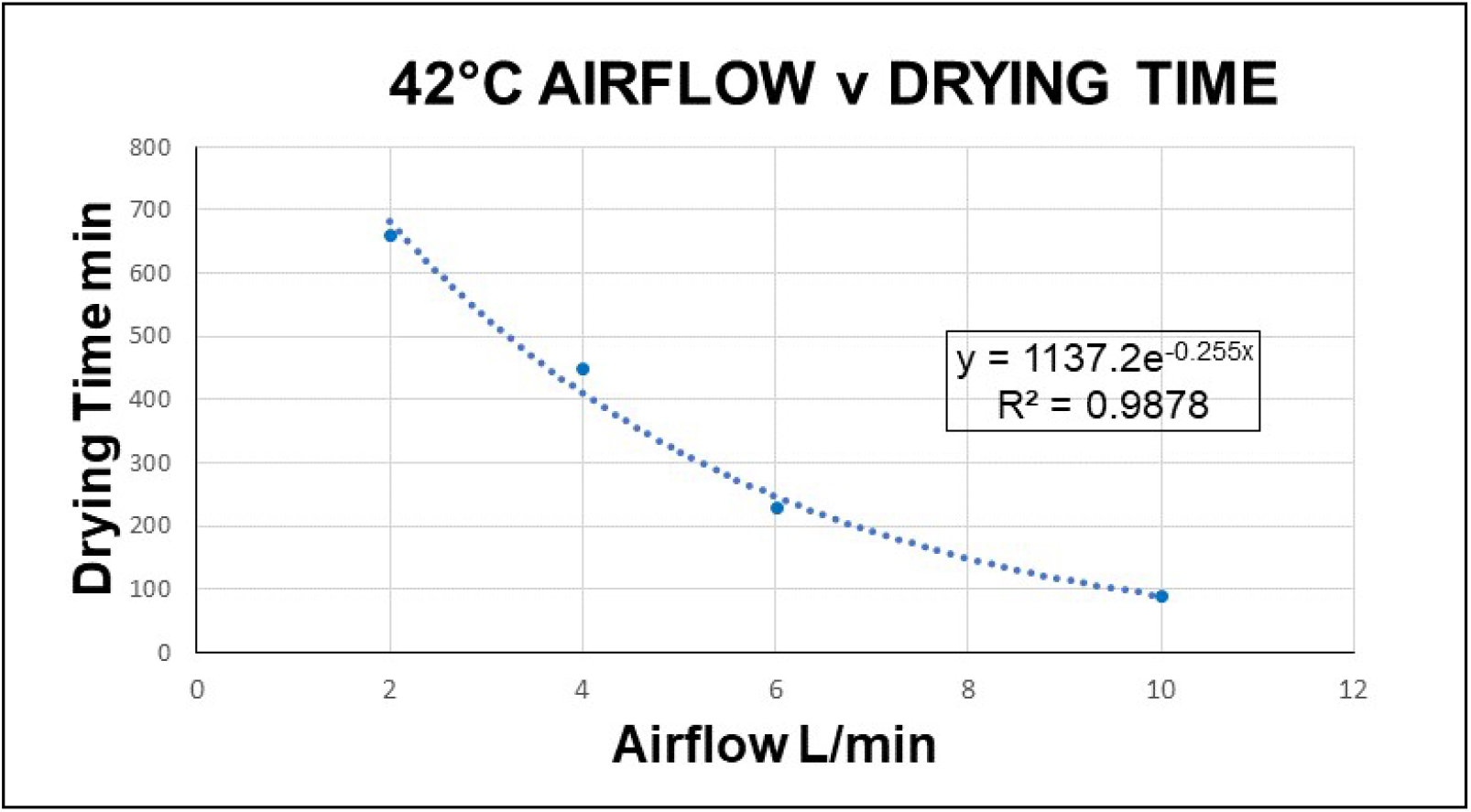
Air flow rate v drying time. Time to achieving plateau dryness shows an exponential relationship versus air flow rate from 2 to 10 L/min, taking from nearly 11 hr to ~1.5 hr. A profile of drying curves at 4 L/min is shown in Figure 3

**Figure 3.**
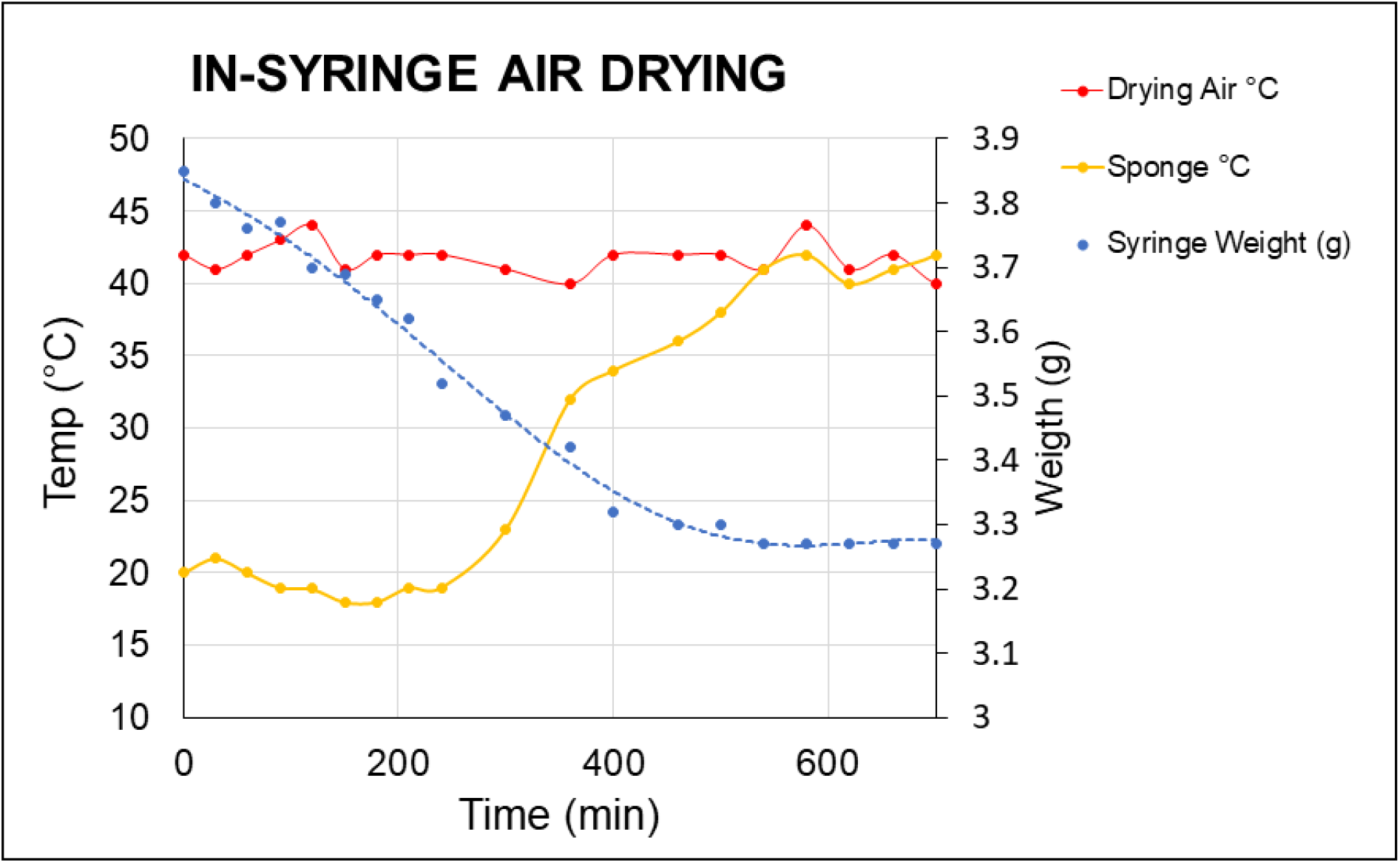
Temperature and weight profiles of drying sponge. Even though exposed to a 4 L/min flow of air at 42°C, the sponge remained at room temperature (18-22°C) for about 4 hr indicating efficient evaporative cooling. This was supported by the progressive loss of weight in the sponge during this period. Sponge temperature began to rise, due to a reduction in evaporation rate as it approached dryness, eventually equalling the inlet air temperature which signalled completion of the drying process, confirmed by no further weight loss.

These data were obtained in single syringe experiments and are not from drying in the rig itself. However, the efficacy of drying in the rig was confirmed by measurement of the very low residual moisture in multiple samples from later drying runs used to produce the batches for further studies.

#### Residual moisture

Analysis by the Karl Fisher method indicated that each STVX-01 sponge contained an average 0.25 mg of water (0.4% w/w), exceeding the acceptance criterion for this type of dried product, which is generally accepted to be less than 3% residual moisture.

#### Recovery of vaccine antigens and adjuvant on rehydration

It was found that near complete delivery of the vaccine from STVX-01 after rehydration required a sustained 30-60 second flicking action to agitate the sponge in the diluent, releasing adherent air bubbles, and fully saturating the sponge.

#### Aluminium hydroxide adjuvant

Full recovery of the adjuvant is essential for the validity of the sponge concept since many of the commonest childhood vaccines are only efficacious when they contain antigens physiochemically bound to adjuvant [21]. Efficacy would be compromised if significant adjuvant damage or entrapment of adjuvant particles occurred in the dried sponges. Absorbance at 960nm was found to be a rapid and simple spectrophotometric method to assess particulate Al(OH)_3_. It was unaffected by trehalose or polysorbate 80 additives and was linear (R2)>0.99 in the tested range of 0.02 – 2.5 mg/ml.

Adjuvant content was also verified using the European pharmacopoeia chemical method for Al^+++^ quantification. The batch of bulk Tetadif liquid vaccine used to produce the three batches of STVX-01 in this experiment has a concentration of 1.22mg/ml of Al(OH)_3_ according to the Certificate of Analysis.

#### Preservation of Adjuvant gel structure

Damage to Al(OH)_3_ adjuvanted vaccines by freezing or drying is manifest by clumping and rapid sedimentation of the adjuvant particles and was readily demonstrated in 1 x G sedimentation columns **Figure 5**.

**Figure 4:**
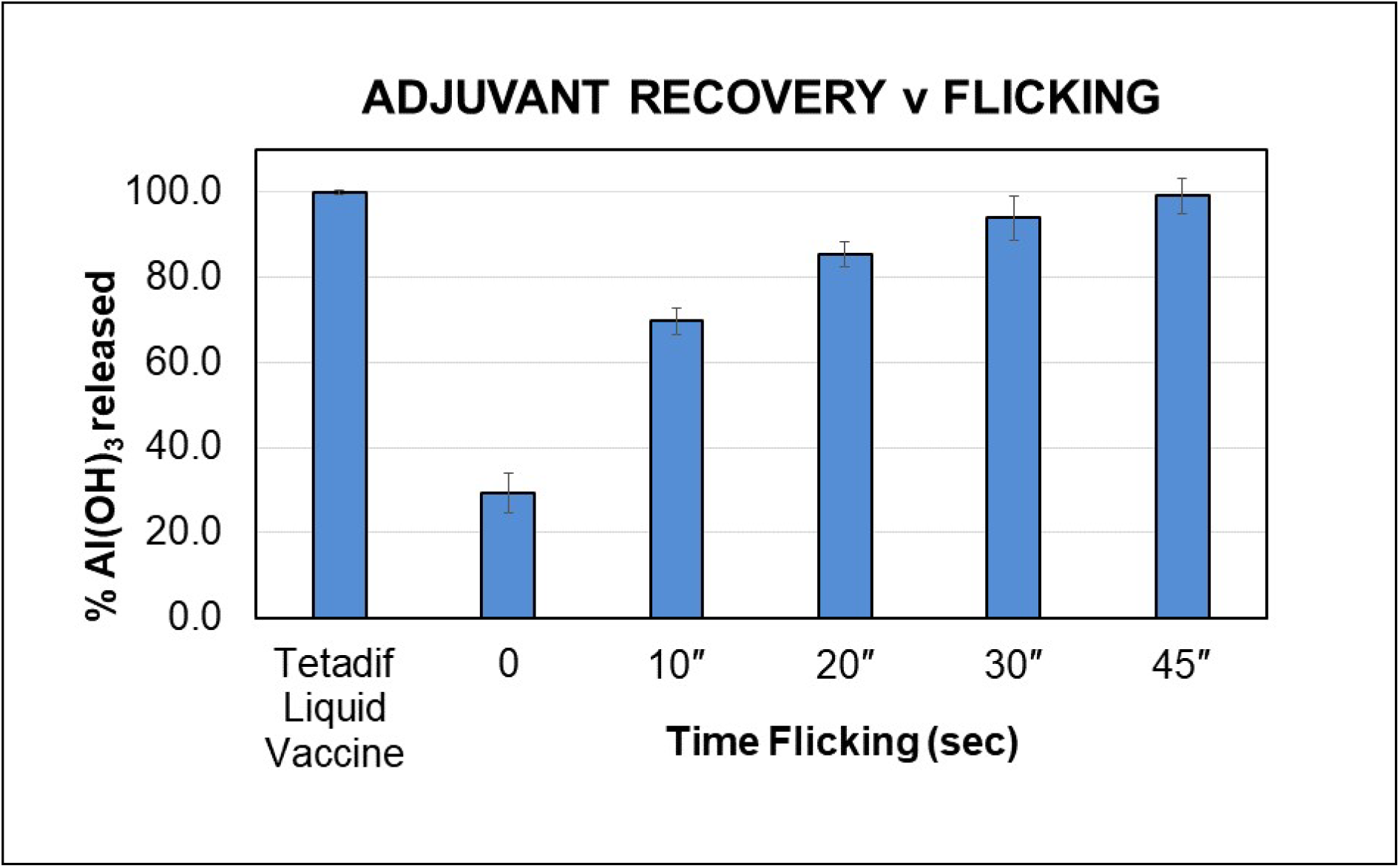
Al(OH)3 release determined by ABS_960_. Complete rehydration and elution of the vaccine components from the sponge requires that water passes through all the pores of the sponge. We found that prolonging the flicking action that is traditionally used to dislodge air bubbles in the syringe prior to injection, was effective in achieving this. Yield progressively increased with duration and 45 seconds was required to ensure full adjuvant release. If flicking was not done, the Al(OH)3 yield obtained was only 29.4%. This was not increased significantly by merely waiting for 45 sec indicating that mechanical agitation was needed (data not shown).

**Figure 5.**
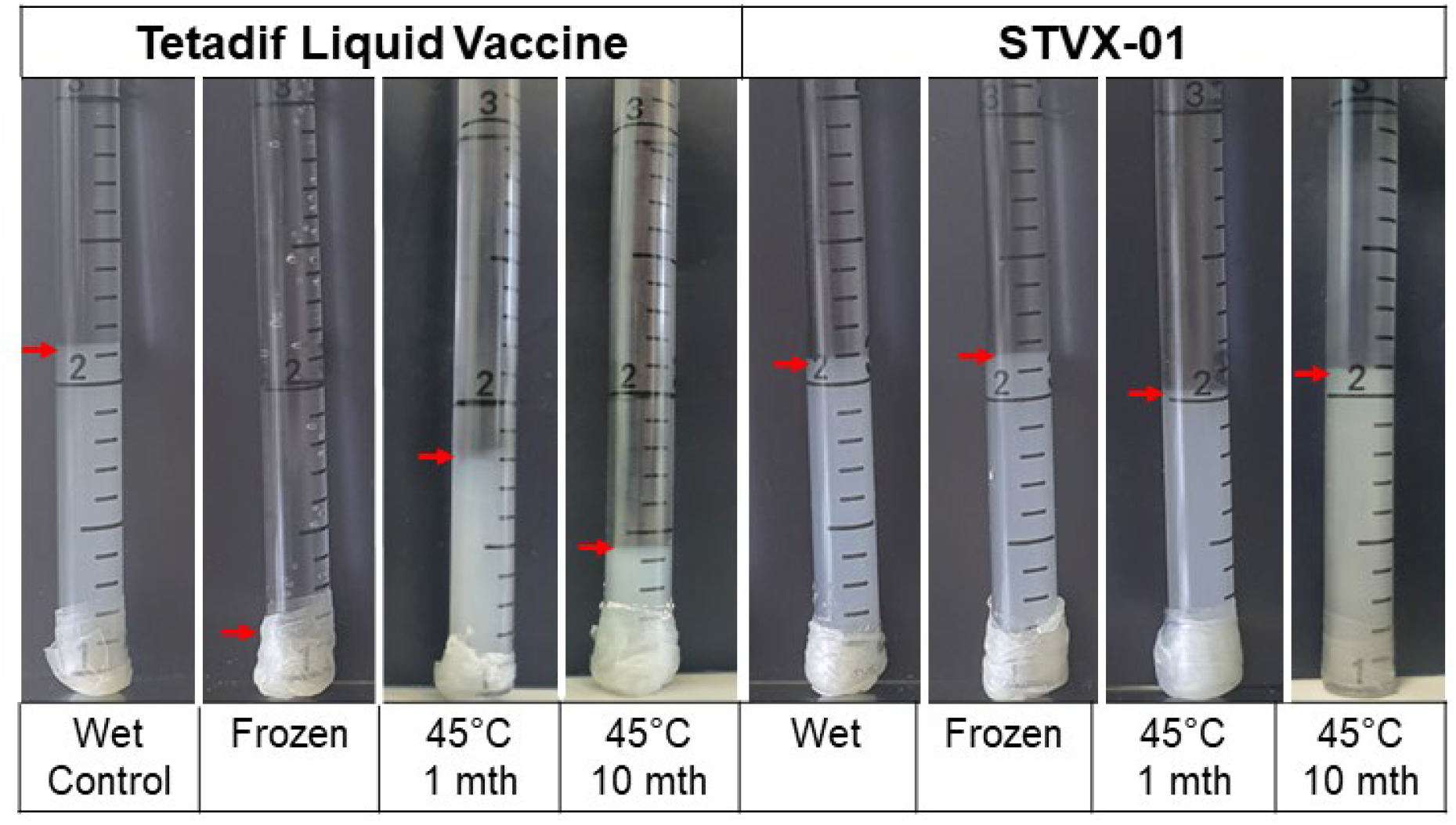
The Tetadif liquid vaccine collapsed and sedimented after freezing or storage at elevated temperature, while the STVX-01 vaccine did not. The disruption of the finely divided sub-micron gel structure of the adjuvant and clumping of the particles was also confirmed by Dynamic Light Scattering (DLS) Figure 6.

**Figure 6.**
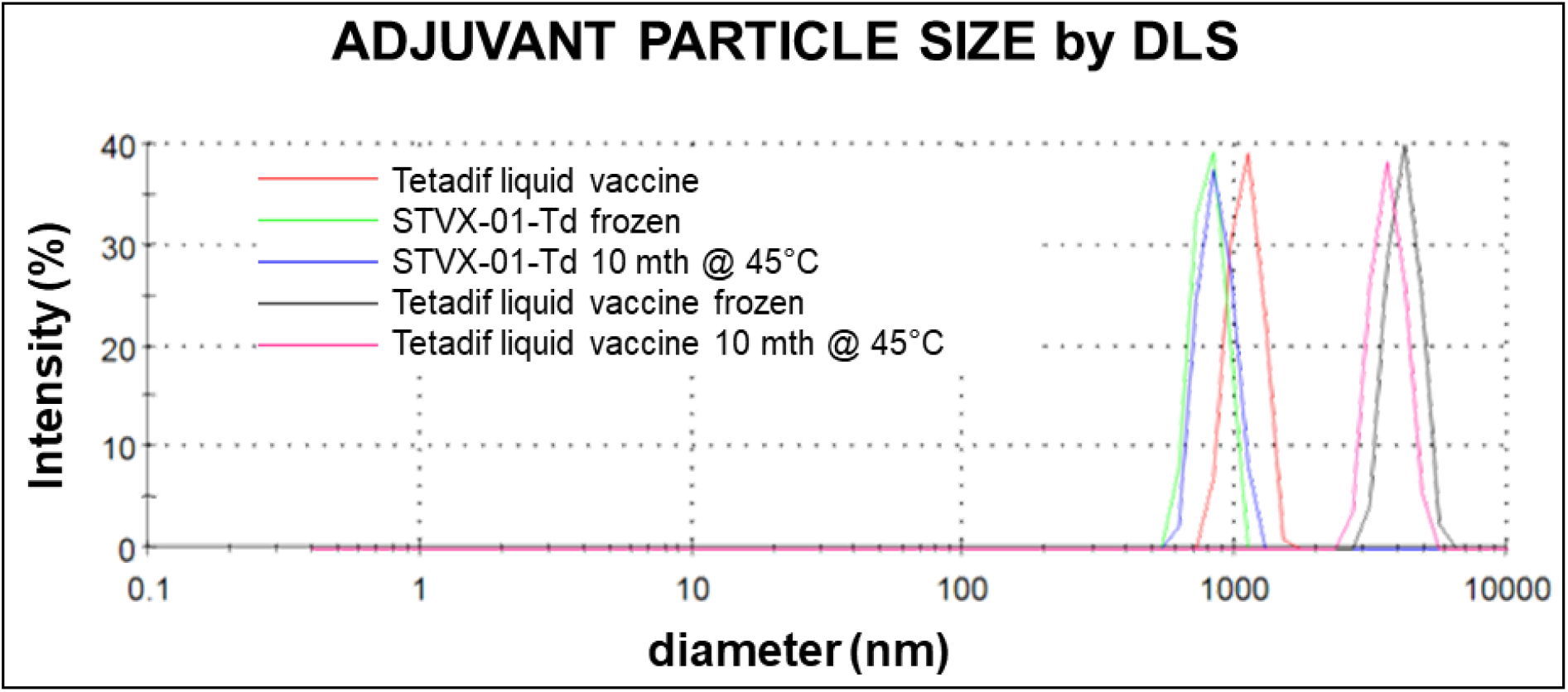
Adjuvant particle size by Dynamic Light Scattering. Particle size of adjuvant in Tetadif liquid vaccine was slightly over 1μm and STVX-01 vaccines either frozen or heated remained submicron. Frozen or heated Tetadif liquid vaccines were aggregated into 3-4 μm diameter clumps.

**Figure 7:**
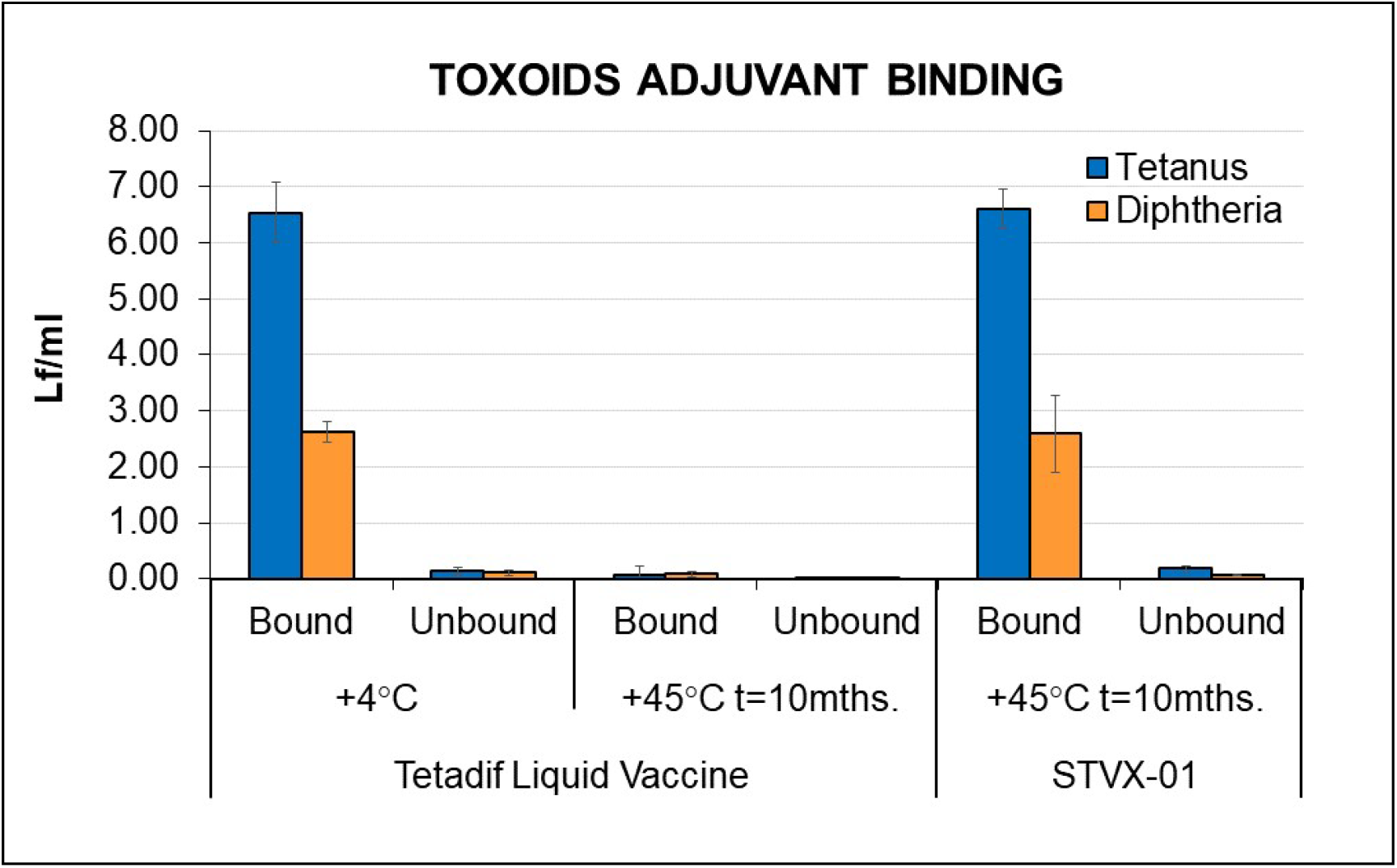
Preservation of bound fraction. Measurements of the residual toxoid in the supernatant after sedimentation of the adjuvant by centrifugation indicated a high level of binding to the adjuvant for both T (98%) and d (96%). Storage of the Tetadif liquid vaccine control at 45°C for 7 months resulted in the antigens progressively degrading at 1 and 7 months, by when they were virtually undetectable either bound T (<2%) or unbound (<2%). In contrast, STVX-01stored for 7 months at 45°C remained intact and fully bound T (97%) and d (97%).

**Figure 8.**
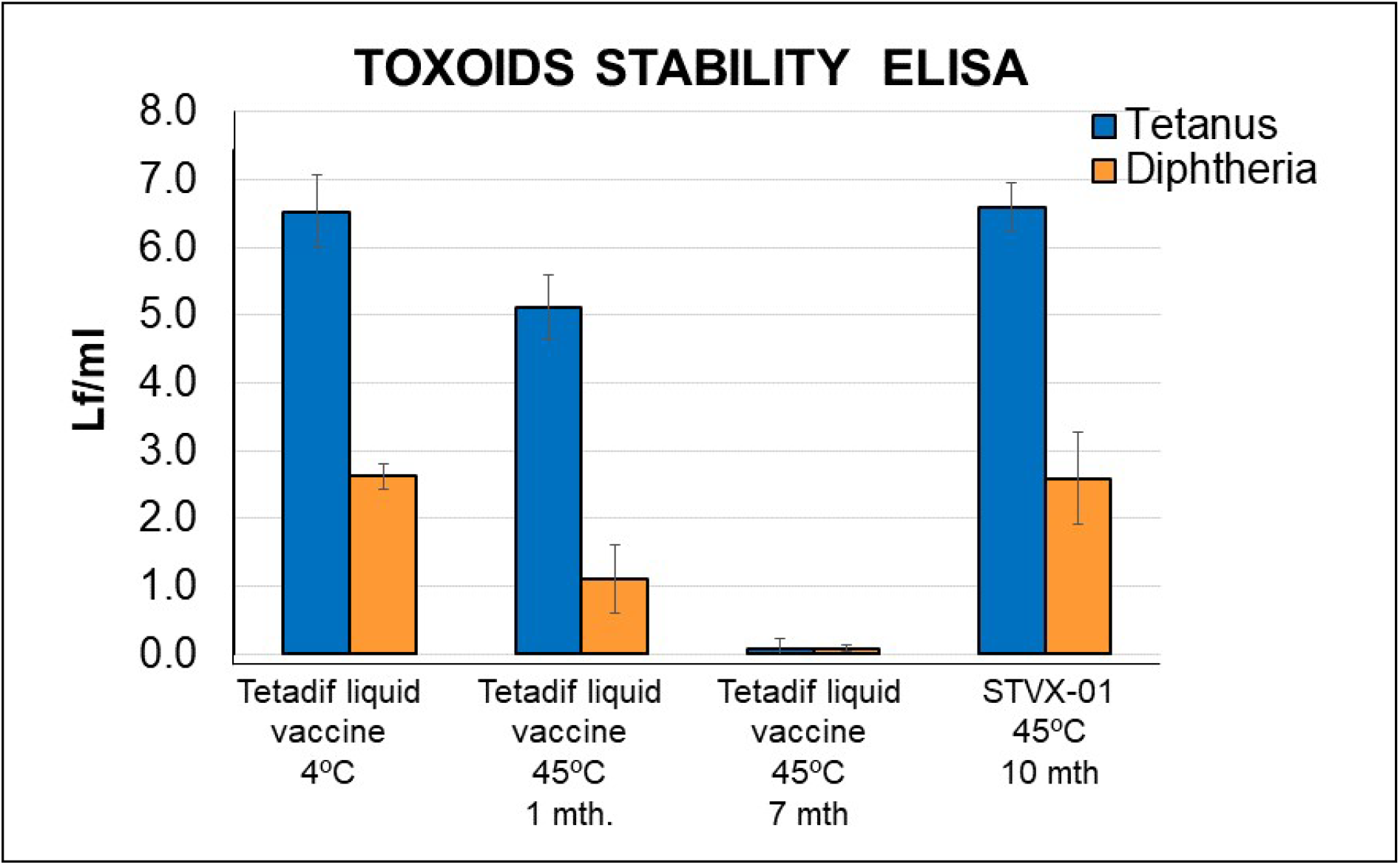
Antigenic integrity STVX-01 after 10 months. ELISA assays showed that the toxoids in Tetadif liquid vaccine were already damaged by 1 month at 45°C and became undetectable by 7 months. In STVX-01 they remained stable at 45° for 10 months.

#### Adjuvant binding

The functional activity of the adjuvant in binding the antigens was determined by measuring toxoid concentrations in the samples by ELISA before and after centrifugation of the adjuvant out of the sample.

#### Antigenic Integrity

The preservation of toxoid antigenicity was measure by a 2-site capture ELISA on fresh, stored and stabilised vaccine.

#### In Vivo Potency of stabilised vaccine

To confirm that the stabilised vaccine was not only intact in the *in vitro* ELISA assay but also retained its immunogenic potency *in vivo*, STVX-01 vaccine after storage at 45°C for 7 and 10 months was compared with the Tetadif liquid vaccine in the Protection Assay routinely used by Bulbio for batch release of their commercial vaccine (Table II)

**Table I.** Measured Al^+++^ concentrations. The Tetadif liquid vaccine and three pools of the eluates of 12 units of STVX-01 were compared. Similar values were obtained for the Tetadif liquid vaccine and the STVX-01 eluates, 1.233 mg/ml ± 0.008 mg/ml for the Tetadif liquid vaccine 1.236 mg/ml ± 0.005 mg/ml for STVX-01.

| Sample | Pooled sample ml | | | $\text{Al}(\text{OH})_3$ mg/ml | | | Average $\text{Al}(\text{OH})_3$ mg/ml |
| --- | --- | --- | --- | --- | --- | --- | --- |
|  | #1 | #2 | #3 | #1 | #2 | #3 |  |
| Tetadif liquid vaccine | 5 | 5 | 5 | 1.22 | 1.24 | 1.23 | 1.233 |
| STVX-01 | 5 | 5 | 5 | 1.24 | 1.23 | 1.24 | 1.236 |

**Table II.** Guinea pigs immunised with either the fresh vaccine or the stabilised vaccine survived the challenge with active toxins, showing that the stabilised vaccine fully retained its function of providing protective immunity against Tetanus and Diphtheria. The retention of full protective functionality in the STVX-01 vaccines after 7-10 months storage at 45°C complies with the WHO guidelines for room temperature Pharmaceuticals [25] no degradation after an accelerated ageing study of 6 months storage at 40°C and 75% RH [26], justifying the title “Fridge Free” vaccines.

| **Tetanus toxin challenge test** N° of survivors/N° of animals challenged* |  |  |  |  |
| --- | --- | --- | --- | --- |
| Number of units /Dilution | Reference Tetanus | Tetadif liquid vaccine | STVX-01 T=0 | STVX-01 T=10mth @45°C |
| 16 | 8/8 | 8/8 | 8/8 | 8/8 |
| 8 | 6/8 | 6/8 | 6/8 | 7/8 |
| 4 | 2/8 | 3/8 | 5/8 | 2/8 |
| 2 | 0/8 | 0/8 | 0/8 | 1/8 |
| POTENCY |  | **86.69 IU**<br>95% CL (55.24-136.18) | **91.72 IU**<br>95% CL (55.07-154.2) | **97.38 IU**<br>95% CL (59.7-159.97) |
| **Diphtheria toxin challenge test** N° of survivors/N° of animals challenged |  |  |  |  |
| Number of units /Dilution | Reference Diphtheria | Tetadif liquid vaccine | STVX-01 T=0 | STVX-01 T=7mth @45°C |
| 16 | 8/8 | 8/8 | 8/8 | 8/8 |
| 8 | 5/8 | 5/8 | 6/8 | 7/8 |
| 4 | 3/8 | 2/8 | 5/8 | 6/8 |
| 2 | 0/8 | 1/8 | 1/8 | 1/8 |
| POTENCY |  | **10.26 IU**<br>95% CL (6.08-17.37) | **14.03 IU**<br>95% CL (8.55-23.49) | **15.00 IU**<br>95% CL (9.47-25.43) |

## DISCUSSION

In the StablevaX process, a porous freeze-dried plug is replaced with a porous air-dried sponge which allows loading the liquid vaccine in a widely dispersed and rapidly drying distribution within the large surface area of the pores of the sponge. This accelerates water evaporation and allows rapid warm-air drying. Filled sponges can be dried thoroughly in 4 - 10 hr depending on the drying temperature and rate of airflow. This provides a fast and economical manufacturing process.

It has been a specific goal of many organisations responsible for delivering vaccines worldwide such as the World Health Organisation (WHO) [27] Medecins Sans Frontieres [28], and Coalition for Epidemic Preparedness Innovations [8] to have vaccines available that do not need refrigeration in a cold chain. The process for making pharmaceuticals and vaccines stable at high and freezing temperatures has been available for some years now,but the preparations on offer, including Hydris (porous membranes in cassettes) [29] microneedle array patches [30], Two chamber syringes [31] etc. have been deemed unsuitable for the rapid and widespread deployment of many vaccines, seemingly on the basis of expense, limited volume capacity, complexity or inconvenience.

We have addressed this problem by developing a thermally stable formulation of vaccine, a full dose of which is absorbed and dried in a sponge. This technique disperses the liquid vaccine widely in the pores of the sponge and ensures simple manufacturing, rapid drying, and rapid reconstitution. The dried sponge is conveniently housed in the barrel of the vaccination syringe itself. It is suitable for most vaccines and, Importantly, adds little to the cost of vaccine delivery in the field and reduces the amount of equipment required. The six or seven separate pieces of equipment required for delivering conventional liquid vaccines in the field (vaccine vial, ± diluent, empty syringe, 2 needles, refrigerator or cold box, certified vaccine thermometer) are replaced by just two, the StablevaX syringe containing the stable vaccine and water for injection. Pre-dosed stable vaccines in the injection syringe guarantees correct dosing and improves safety. It also eliminates the estimated 50% wastage of thermally unstable vaccines [2,3,5,8]

A programme of selection for the properties of the optimal sponge was undertaken and samples of 50 porous matrices, of various types and made from various materials were obtained from multiple manufacturers worldwide, and screened. Some were rejected because they were non-absorbent closed-cell foams, made from non-approvable or toxic materials, of inappropriate pore sizes, or for other technical reasons. The types of open cell matrices we identified as possessing most of the required properties were cellulose foams, reticulated melamine formaldehyde foams and reticulated polyurethane, (both polyether and polyester) foams.

A comparison of these showed that certain polyurethane foams were preferred in that they were, free of plasticisers or other toxic additives, were non-swelling, physically strong and amenable to vigorous, thorough washing and sterilising, and were inexpensive. Also, some were already approved for prolonged exposure to sub-cutaneous tissues as contact dressings for open wounds which testified to their nontoxic and non-irritant status. Some otherwise suitable sponges were not very hydrophilic and absorbed water slowly. This could be overcome by the addition of small quantities of an approved surfactant, and both the initial vaccine loading, and the subsequent elution were greatly improved.

A variety of loading and drying technologies for the sponges were examined before arriving at the “in syringe” process eventually used. It was deemed easier and safer to preserve sterility by loading and drying the sterile sponges already loaded into sterile syringes. This advantage is, of course, balanced by the batch nature of this technique and the possible difficulty and complexity of scaling up to millions of StablevaX syringes. Alternative processes more suited to high volume production are being investigated. A modest temperature was initially chosen for the drying air to minimise any potential product damage. Since the sponge reached a low plateau weight at the end of the drying experiments, the product was assumed to have reached its minimum moisture content at this stage. This was confirmed by showing a very low residual moisture content by Karl Fisher analysis.

It is clear from these data that faster drying could probably be achieved without damaging the product by using higher temperature drying air in an industrial process. It is not clear from these experiments whether the low temperature maintained initially in the sponges was due to evaporative cooling, or due to the thermistor being located towards the base of the sponge which obstructed airflow and protected it from heat until the drying front of the vaccine reached it. Whatever the case, drying to a very low residual moisture content was achieved in a reasonable time and produced suitable batches for the further studies.

It is worth noting that the amount of pharmaceutical material stored and delivered by this method can vary over a very wide range by tailoring the size of the sponge insert and the size of syringe required, to fit any dose. Since the sponge is ~90% void volume and chosen to have a very high capacity to hold aqueous products (of the order of 25 millilitres per gram), and is not swollen significantly by the absorbate, there is no need to increase the size of the syringe to allow for the loading of product solution into the sponge insert. Theoretically, there is no practical limit to the size of the syringe (or other container), or of the sponge.

Unlike some alternative techniques to produce fridge-free vaccines such as micro array needle patches [32] or membrane-based cassettes [33], this method is not restricted to small volumes of potent substances. The full 0.5 ml dose of current vaccines can be accommodated by the sponge and successfully stabilised obviating any need for a separate concentration step.

It has the additional advantage of using syringe-based injection, a technique in which current vaccinators are trained and competent, and it gives assurance that the correct dose of vaccine has indeed been injected into every patient.

In summary, a commercial Aluminium Hydroxide adjuvanted Tetanus/diphtheria booster vaccine, was fully functional and protective *in vivo* against lethal toxin challenge after storage in this device at 45°C for at least 7 months. This confirms that the StablevaX technology is a feasible technique that permits fridge-free storage of potent vaccines.

## CONCLUSIONS

A fridge-free, single-dose, thermostable vaccine product has been developed in the plastic syringe used for vaccination. The vaccine is stabilised using the inert disaccharide trehalose as a sugar glass in the pores of a compressible sponge. Both the aluminium hydroxide adjuvant and the tetanus and diphtheria toxoids of the divalent Td vaccine used were shown to be fully recoverable with their physicochemical properties intact, after storage for many months at 45°C. Since trehalose has been approved as a stabiliser in other marketed pharmaceuticals, a wide range of other vaccines could potentially benefit from this process and the mandatory use of the cold chain for these would not be necessary. The Td vaccine was fully potent in batch release protection studies against supra-lethal doses of active tetanus and diphtheria toxins. These assays had been commercially developed by the vaccine manufacturer to assure that manufactured vaccine batches could be released for sale as being fully protective. It is therefore very likely that the StablevaX process will prove to be a safe and effective process for thermostabilising vaccines for clinical use in humans. The widespread adoption of the StablevaX process would facilitate the distribution of many lifesaving medicines to the countries lacking secure refrigeration infrastructure.

## ACKNOWLEDGEMENTS

We are grateful to BulBio (BB – NCIPD Ltd. Sofia. Bulgaria) who provided the commercial Td vaccine Tetadif™ and performed the protective potency studies in Guinea pigs, to Stablepharma Spain Ltd. (Madrid, Spain) for the laboratory experiments, provision of assistance and facilities, and to Stablepharma Ltd for advice and financial support. We would like to thank staff of the Biological Services Division and Standards Processing Division at the National Institute for Biological Standards and Control (NIBSC South Mimms. London) for their advice and technical contributions to this work.

## BIBLIOGRAPHY

[1] WHO 2019 NCoV UCC Systems Pfizer BioNTech Vaccine 2022.1 Eng | PDF, (n.d.). https://es.scribd.com/document/559688399/WHO-2019-NCoV-UCC-Systems-Pfizer-BioNTech-Vaccine-2022-1-Eng (accessed August 31, 2022).

[2] Monitoring vaccine wastage at country level Guidelines for programme managers Immunization, Vaccines and Biologicals, 2005. https://apps.who.int/iris/handle/10665/68463 (accessed September 8, 2022).

[3] V.C. Falcón, Y.V.V. Porras, C.M.G. Altamirano, U. Kartoglu, A vaccine cold chain temperature monitoring study in the United Mexican States, Vaccine. 38 (2020) 5202–5211. https://doi.org/10.1016/j.vaccine.2020.06.014.

[4] D. Kristensen, D. Chen, R. Cummings, Vaccine stabilization: Research, commercialization, and potential impact, Vaccine. 29 (2011) 7122–7124. https://doi.org/10.1016/j.vaccine.2011.05.070.

[5] D.M. Matthias, J. Robertson, M.M. Garrison, S. Newland, C. Nelson, Freezing temperatures in the vaccine cold chain: A systematic literature review, Vaccine. 25 (2007) 3980–3986. https://doi.org/10.1016/j.vaccine.2007.02.052.

[6] D. McAdams, D. Chen, D. Kristensen, Spray drying and vaccine stabilization., Expert Rev Vaccines. 11 (2012) 1211–1219. https://doi.org/10.1586/erv.12.101.

[7] A. Sharma, D. Khamar, S. Cullen, A. Hayden, H. Hughes, Innovative Drying Technologies for Biopharmaceuticals, Int J Pharm. 609 (2021). https://doi.org/10.1016/j.ijpharm.2021.121115.

[8] G. Kanojia, R. ten Have, P.C. Soema, H. Frijlink, J.P. Amorij, G. Kersten, Developments in the formulation and delivery of spray dried vaccines, Hum Vaccin Immunother. 13 (2017) 2364–2378. https://doi.org/10.1080/21645515.2017.1356952.

[9] Roser Bruce, Injections, US10821210B2 n.d., (n.d.). https://worldwide.espacenet.com/patent/search/family/045876206/publication/US10821210B2?q=US10821210B2.

[10] K.R. Ward, G.D.J. Adams, H.O. Alpar, W.J. Irwin, Protection of the enzyme L-asparaginase during lyophilisation-a molecular modelling approach to predict required level of lyoprotectant, 1999. https://doi.org/10.1016/s0378-5173(99)00163-5.

[11] Roser Bruce, Garcia de Castro Arcadio, WO1999047174A1 Amorphous Glasses for Stabilizing Sensitive products, (n.d.). https://worldwide.espacenet.com/patent/search/family/026313295/publication/WO9947174A1?q=WO9947174A1.

[12] Roser Bruce, Colaco Camilo, WO9118091A1 STABILISATION OF BIOLOGICAL MACROMOLECULAR SUBSTANCES AND OTHER ORGANIC COMPOUNDS, n.d., (n.d.). https://worldwide.espacenet.com/patent/search/family/010675945/publication/WO9118091A1?q=WO9118091. (accessed September 7, 2022).

[13] N.K. Jain, I. Roy, Effect of trehalose on protein structure, Protein Science. 18 (2009) 24–36. https://doi.org/10.1002/pro.3.

[14] Roser Bruce, GB2187191A protection of proteins and the like, (n.d.). https://worldwide.espacenet.com/patent/search/family/027449677/publication/GB2187191A?q=GB2187191A.

[15] Roser Bruce, Colaco Camilo, Kampinga Jaap, Smith Christopher, US6890512B2 Methods of preventing aggregation of various substances upon rehydration or thawing and compositions obtained thereby, n.d., (n.d.). https://worldwide.espacenet.com/patent/search/family/022958303/publication/US6890512B2?q=US6890512B2 (accessed September 7, 2022).

[16] J L Green, C A Angel, Phase relations and vitrification in saccharide-water solutions and the trehalose anomaly, 1989. https://doi.org/10.1021/J100345A006.

[17] X. Tang, M.J. Pikal, Design of Freeze-Drying Processes for Pharmaceuticals: Practical Advice, 2004. https://doi.org/10.1023/B:PHAM.0000016234.73023.75.

[18] Bul Bio-National Center of Infectious and Parasitic Diseases Ltd., tetadif, (n.d.). https://extranet.who.int/pqweb/content/tetadif (accessed September 7, 2022).

[19] N. Dumpa, K. Goel, Y. Guo, H. McFall, A.R. Pillai, A. Shukla, M.A. Repka, S.N. Murthy, Stability of Vaccines, AAPS PharmSciTech. 20 (2019). https://doi.org/10.1208/s12249-018-1254-2.

[20] S.G. Reed, M.T. Orr, C.B. Fox, Key roles of adjuvants in modern vaccines, Nat Med. 19 (2013) 1597–1608. https://doi.org/10.1038/nm.3409.

[21] L. Wolff, J. Flemming, R. Schmitz, K. Gröger, C. Müller-Goymann, Protection of aluminum hydroxide during lyophilisation as an adjuvant for freeze-dried vaccines, Colloids Surf A Physicochem Eng Asp. 330 (2008) 116–126. https://doi.org/10.1016/j.colsurfa.2008.07.031.

[22] X. Lai, Y. Zheng, I. Søndergaard, H. Josephsen, H. Løwenstein, J.N. Larsen, H. Ipsen, S. Jacobsen, Determination of aluminium content in aluminium hydroxide formulation by FT-NIR transmittance spectroscopy, Vaccine. 25 (2007) 8732–8740. https://doi.org/10.1016/j.vaccine.2007.10.026.

[23] L. Coombes, P. Stickings, R. Tierney, P. Rigsby, D. Sesardic, Development and use of a novel in vitro assay for testing of diphtheria toxoid in combination vaccines, J Immunol Methods. 350 (2009) 142–149. https://doi.org/10.1016/j.jim.2009.09.002.

[24] L. Coombes, R. Tierney, P. Rigsby, D. Sesardic, P. Stickings, In vitro antigen ELISA for quality control of tetanus vaccines, Biologicals. 40 (2012) 466–472. https://doi.org/10.1016/j.biologicals.2012.07.011.

[25] WHO guidelines on stability testing of active pharmaceutical ingredients and finished pharmaceutical products, (n.d.). https://www.who.int/publications/m/item/who-guidelines-on-stability-testing-of-active-pharmaceutical-ingredients-and-finished-pharmaceutical-products (accessed October 7, 2022).

[26] No. 1010, 2018 WHO Technical Report Series, Stability testing of active pharmaceutical ingredients and finished pharmaceutical products, n.d. https://www.who.int/publications/m/item/who-guidelines-on-stability-testing-of-active-pharmaceutical-ingredients-and-finished-pharmaceutical-products.

[27] U.J. Lloyd, Technologies for vaccine delivery in the 21st century. World Health Organization Geneva 2000 in collaboration with DEPARTMENT OF VACCINES AND BIOLOGICALS, 2000. https://apps.who.int/iris/bitstream/handle/10665/66569/WHO_VB_00.35-eng.pdf?sequence=1.

[28] Heat-stable vaccines urgently needed to reach the one in five children missed by immunisation worldwide | MSF, (n.d.). https://www.msf.org/heat-stable-vaccines-urgently-needed-reach-one-five-children-missed-immunisation-worldwide.

[29] R. Alcock, M.G. Cottingham, C.S. Rollier, J. Furze, S.D. de Costa, M. Hanlon, A.J. Spencer, J.D. Honeycutt, D.H. Wyllie, S.C. Gilbert, M. Bregu, A.V.S. Hill, Long-term thermostabilization of live poxviral and adenoviral vaccine vectors at supraphysiological temperatures in carbohydrate glass, Sci Transl Med. 2 (2010). https://doi.org/10.1126/scitranslmed.3000490.

[30] F.S. Quan, Y.C. Kim, D.G. Yoo, R.W. Compans, M.R. Prausnitz, S.M. Kang, Stabilization of influenza vaccine enhances protection by microneedle delivery in the mouse skin, PLoS One. 4 (2009). https://doi.org/10.1371/journal.pone.0007152.

[31] Primary Packaging | Vetter, (n.d.). https://www.vetter-pharma.com/en/services/packaging-and-assembly/primary-packaging/ (accessed September 1, 2022).

[32] J. Li, M. Zeng, H. Shan, C. Tong, Microneedle Patches as Drug and Vaccine Delivery Platform, Curr Med Chem. 24 (2017). https://doi.org/10.2174/0929867324666170526124053.

[33] HydRIS (Hypodermic Rehydration Injection System), (n.d.). https://www.novalabs.co.uk/stabilisation-technologies.

